# Analysis of The Senescence Secretome During Zebrafish Retina Regeneration

**DOI:** 10.1101/2025.01.31.635944

**Authors:** Gregory J. Konar, Kyle T. Vallone, Tu D. Nguyen, James G. Patton

**Author notes:** Authors contributed equally to this work (Gregory J. Konar, Kyle T. Vallone). **Corresponding author:** James G Patton, PhD, Department of Biological Sciences, Vanderbilt University.

## Abstract

Zebrafish possess the innate ability to regenerate any lost or damaged retinal cell type with Müller glia serving as resident stem cells. Recently, we discovered that this process is aided by a population of damage-induced senescent immune cells. As part of the Senescence Associated Secretory Phenotype (SASP), senescent cells secrete numerous factors that can play a role in the modulation of inflammation and remodeling of the retinal microenvironment during regeneration. However, the identity of specific SASP factors that drive initiation and progression of retina regeneration remains unclear. Here, we mined the SASP Atlas and RNAseq datasets to identify differentially expressed SASP factors after retina injury, including two distinct acute damage regimens, as well as a chronic, genetic model of retina degeneration. We discovered a 31-factor “Regeneration-associated Senescence Signature” (RASS) that represents SASP factors and senescence markers that are conserved across all data sets and are upregulated after damage. Among these, we show that depletion of Nucleophosmin 1 (*npm1a)* inhibits retina regeneration. Our data support the model that differential expression of SASP factors promotes regeneration after both acute and chronic retinal damage.

## Introduction

The retina is a complex tissue within the central nervous system that controls our ability to see and perceive the world around us. It comprises photoreceptors (rods and cones), interneurons (bipolar, horizontal, and amacrine cells), retinal ganglion cells, and glial cells including Müller glia (MG) and microglia/macrophages (Marc 1998, Masland 2001). After acute injury in mammals, whether from stress, toxic exposure, or trauma, retinal neurons die and are lost permanently. However, in certain species, including the teleost fish *Danio rerio*, MG can respond to acute damage and produce progenitor cells that migrate to the site of damage and replace any lost cell type, successfully allowing for regeneration of damaged neurons and restoration of lost vision (Yurco and Cameron 2005, Fausett and Goldman 2006, Raymond, Barthel et al. 2006, Lenkowski and Raymond 2014, Wan and Goldman 2016, Konar, Ferguson et al. 2020). Studies have been conducted using single-cell RNA sequencing (scRNAseq) and bulk RNAseq on retinas from mammals, birds, and zebrafish for comparative analysis of gene expression patterns, resulting in the identification of regulatory networks that promote or inhibit retina regeneration (Hoang, Wang et al. 2020, Palazzo, Todd et al. 2022). The collective data illustrate clear differences in how MG respond to damage and highlight the role that inflammation and microglia/macrophages play in chronic and acute damage models (Mitchell, Lovel et al. 2018, Silva, Nagashima et al. 2020, Nagashima and Hitchcock 2021, Iribarne and Hyde 2022, Zhou, Zhang et al. 2022, Bludau, Weber et al. 2024)

Historically, senescence has been thought to be mostly involved in aging and age-associated diseases, but has more recently been shown to paradoxically play a role in wound healing and regeneration (Andrade, Sun et al. 2022, Ring, Valdivieso et al. 2022, Kita, Yamamoto et al. 2024). Senescence occurs when a cell irreversibly withdraws from the cell cycle, due to the end of replicative lifespan or a variety of cellular stresses (Rodier and Campisi 2011, Campisi 2013, Dodig, Čepelak et al. 2019). These stresses are highly heterogenous in nature and include aging, telomere shortening, oxidizing agents, or DNA damaging agents (Bitencourt, Vargas et al. 2024, Xu, Cao et al. 2024). Senescence can occur in all somatic cells, including both proliferation-competent and post-mitotic cells, and in cancer cells (Huang, Hickson et al. 2022). In many degenerative diseases, senescent cells accumulate and disrupt cellular processes, allowing for disease progression and decline of tissue function (He and Sharpless 2017, Adams, Schupp et al. 2020, Kirkland and Tchkonia 2020, Gasek, Kuchel et al. 2021).

Even though senescent cells exit the cell cycle, they remain metabolically active, undergo significant chromatin remodeling events, and release numerous signaling factors as part of the Senescence Associated Secretory Phenotype (SASP)(Dodig, Čepelak et al. 2019, Rhinn, Ritschka et al. 2019). While identifying specific markers of senescence is ongoing, expression of *p16, p21, p53*, and *Rb* are upregulated when cells undergo senescence, and senescent cells also upregulate a mitochondrial β-galactosidase that is a commonly used marker of senescence (Lee, Han et al. 2006, Fan, Jiang et al. 2011, Mosteiro, Pantoja et al. 2018, Ritschka, Knauer-Meyer et al. 2020).

The SASP secretome is a collection of secreted factors including pro-inflammatory cytokines, chemokines, growth factors, proteases, and extracellular matrix (ECM) components (Kuilman and Peeper 2009, Coppé, Desprez et al. 2010, Oubaha, Miloudi et al. 2016, Lazzarini, Nicolai et al. 2018). These factors act on the local microenvironment in a paracrine manner to promote cellular processes ranging from inflammation modulation to ECM degradation and remodeling, and cleavage of signaling molecules (Acosta, Banito et al. 2013, Tchkonia, Zhu et al. 2013). SASP factors were originally reported in human fibroblasts via large scale genomic studies, but are released by all senescent cells, including pigmented epithelial cells in the retina (Hernandez-Segura, de Jong et al. 2017, Lazzarini, Nicolai et al. 2018, George, Lu et al. 2021). The SASP secretome is highly heterogeneous, with localized populations of senescent cells showing functional heterogeneity in respect to both their expression of senescence markers and the overall composition of secreted factors (Bitencourt, Vargas et al. 2024). This heterogeneity is also dependent on how senescence is induced, with the age-induced SASP secretome showing considerable variation from the SASP of senescent cells induced via irradiation or other DNA damaging agents (Hernandez-Segura, de Jong et al. 2017, Wechter, Rossi et al. 2023). Multiple databases of SASP factors exist, but of the over 2000 genes observed in senescent cells, only around 5% of them form the “Core Senescence Associated Signature,” with many others being context-dependent additions to the SASP (Basisty, Kale et al. 2020, Saul, Kosinsky et al. 2022, Suryadevara, Hudgins et al. 2024). Thus, understanding the SASP of senescent cells in different contexts is needed to precisely define how senescence exerts both beneficial and harmful effects on the cellular microenvironment during regeneration. Identifying specific factors may provide unique therapeutic targets that could help induce regeneration in organisms where such repair is blocked.

Recently, we showed that damage-induced senescent immune (DISI) cells promote regeneration of retinal neurons after acute damage in zebrafish, consistent with findings that senescent cells can have a pro-regenerative influence in the spinal cord and fin of zebrafish (Da Silva-Alvarez, Guerra-Varela et al. 2020, Paramos-de-Carvalho, Martins et al. 2021, Konar, Flickinger et al. 2024). This pro-regenerative influence is thought to be the result of SASP factors that can influence both the inflammatory microenvironment and stem cell niches that drive regeneration. In the retina, MG are highly sensitive to inflammation, with too much or too little inflammatory signaling being detrimental to regeneration (Ritschka, Storer et al. 2017, Lahne, Nagashima et al. 2020, Iribarne 2021, Leach, Hanovice et al. 2021, Lee, Lin et al. 2021, Nagashima and Hitchcock 2021, Iribarne and Hyde 2022). Like inflammation, the appearance and clearance of senescent cells appears to be required for regeneration. The identity of specific SASP factors during regeneration, how those factors change from initiation to progression of regeneration, and whether there is a regeneration-based SASP signature remain unknown. Here, we analyzed our own and publicly available datasets from different models of retina damage to identify a core group of 31 SASP factors that are differentially expressed during retina regeneration. Among these factors, we find that depletion of the chromatin remodeling factor *npm1a* can inhibit regeneration.

## Materials and Methods

### Zebrafish SASP Factor Identification

To build a database of SASP factors (**Table S1**) for analysis in the regenerating zebrafish retina, we obtained zebrafish orthologs of soluble proteins from human SASP conditions in the SASP Atlas (Basisty, Kale et al. 2020). The UniProt IDs of soluble proteins with a log_2_FC < 2, *p*-value ≤ 0.001, and *q*-value ≤ 0.05 from any of the following senescence conditions were extracted from the SASP Atlas: irradiated renal epithelial cells, irradiated fibroblasts, inducible Ras overexpression (4 and 7 days of induction) in fibroblasts, or atazanavir-treated (9 and 14 days of treatment) fibroblasts (Basisty, Kale et al. 2020). These UniProt IDs were converted to human ENSEMBL gene IDs using the UniProt.ws package (v. 2.46.1) (Carlson 2024) in R (v. 4.4.1)(R Development Core Team 2010). To obtain zebrafish orthologs, human Entrez gene IDs and ZFIN gene IDs from a zebrafish-human ortholog table maintained by ZFIN (accessed October 23, 2024) were converted to their species’ respective ENSEMBL gene IDs using the biomaRt package (v. 2.60.1) (Durinck, Spellman et al. 2009). The resulting tables were matched by human ENSEMBL ID. For any genes that biomaRt were unable to assign a zebrafish ENSEMBL ID, orthologs (when available) were manually added from a search of the ZFIN database, resulting in a total of 744 orthologs identified. Additionally, nine genes known to be markers of senescence were added to this database: *myca, mycb, h2ax, h2ax1, jun*, *cdkn1a, cdkn2a/b, rb1, and tp53* (Dodig, Čepelak et al. 2019, Saul, Kosinsky et al. 2022, Suryadevara, Hudgins et al. 2024).

### RNAseq datasets

We used several public RNAseq datasets from recent publications, representing a variety of damage paradigms, cell types, and durations, to analyze zebrafish retina regeneration through the lens of senescence and the SASP. Primarily, we focused on photoreceptor injury induced via light lesion or chemical ablation of retinal ganglion cells with N-methyl-D-aspartate (NMDA). Additionally, we included a whole retina, bulk RNAseq dataset generated by our lab (GSE277792) to provide timepoints (3 dpi, 12 dpi, 20 dpi) through late retina regeneration for the NMDA damage model. The properties of the RNA sequencing datasets analyzed in this study are summarized in **Table 1**.

**Table 1.** RNA sequencing datasets used in analysis.

| Reference | Data accession | Injury model(s) | Cell types | Time points post injury | Sequencing method |
| --- | --- | --- | --- | --- | --- |
| Konar et al. 2025, *Front Aging* (this publication) | GSE277792 | NMDA treatment | whole retina | 3d, 12d, 20d | paired-end RNA sequencing |
| Hoang et al. 2020, *Science* | GSE135406 | NMDA treatment, acute light lesion | Müller glia (gfap:GFP<sup>+</sup>) and GFP<sup>-</sup> retinal populations | 4h, 10h, 20h, 36h | paired-end RNA sequencing |
| Hoang et al. 2020, *Science* | https://github.com/jiewwwang/Single-cell-retinal-regeneration, https://proteinpaint.stjude.org/F/2019.retina.scRNA.html | NMDA treatment, acute light lesion | whole retina | 4h, 10h, 20h, 36h | single-cell RNA sequencing |
| Kramer et al. 2021, *Front Cell Dev Biol* | GSE180518 | acute light lesion | whole retina | 24h, 36h, 72h, 5d, 10d, 14d, 28d | 3' mRNA sequencing |
| Kramer et al. 2023, *Front Cell Dev Biol* | GSE233896 | chronic light lesion | whole retina | 24h, 36h, 72h, 5d, 10d, 14d, 28d | 3' mRNA sequencing |
| Grabinski et al. 2023, *Front Mol Neurosci* | GSE223277, PRJNA787536 | genetic models of photoreceptor regeneration (cep290 and bbs2 mutants) | whole retina | 6 months post fertilization | paired-end RNA sequencing |

### Long term NMDA damage RNAseq dataset generation

Adult zebrafish were intravitreally injected with 0.5 µL of 10mM NMDA and then sacrificed at 3-, 12-, and 20-days post injection. Each time point was the product of 3 biological replicates that themselves were the product of 3 pooled retinas. Retinas were collected into TRIzol and total RNA was collected using phenol-chloroform extraction. Sample libraries were prepared by the Vanderbilt VANTAGE sequencing core using a stranded mRNA (polyA-selected) library preparation kit (New England Biolabs). Libraries were sequenced on the NovaSeq X Plus by the core facility at a depth of 50M reads per sample, and all raw data were submitted to the NCBI GEO database (GSE277792). Analysis of these datasets involved cross-reference to control undamaged zebrafish (GSE223277) for determination of gene enrichment by NMDA damage (Grabinski, Parsana et al. 2023).

### RNAseq data processing

Raw read files from each bulk RNAseq dataset were imported into the Galaxy web platform (The Galaxy Community 2024). The public server at usegalaxy.org was used to align reads to the *Danio rerio* genome assembly (GRCz11, ENSEMBL release 111) using STAR (v. 2.7.11a) (Dobin, Davis et al. 2013). Alignment results were analyzed using MultiQC (v. 1.24.1) and are available in **Table S2** along with sample metadata (Ewels, Magnusson et al. 2016). For technical replicate pairs in the Hoang et al. (2020) bulk RNAseq datasets, replicates with low numbers of mapped reads were removed prior to analysis (Hoang, Wang et al. 2020). One sample from Kramer et al. (2021) was also excluded from analysis due to a low mapping rate and is notated as such in **Table S2** (Kramer, Gurdziel et al. 2021).

Transcript-level count data were imported into R using the tximport package (v. 1.32.0) and differential gene expression analysis was performed using DESeq2 (v. 1.44.0) (Love, Huber et al. 2014, Soneson 2016). All experimental treatments were compared to the respective naïve controls indicated in **Table S2** rather than dark-adapted controls to facilitate comparisons across the NMDA and light damage paradigms. Principal Component Analysis plots are available for each set of comparisons (**Figure S1)** and the results of the differential gene expression analysis for all comparisons are available as **Table S3**. Volcano plots and heatmaps of the data were generated using ggplot2 (v. 3.5.1) and ComplexHeatmap (v. 2.22.0), respectively (Gu, Eils et al. 2016, Wickham 2016, Gu 2022).

Seurat (v. 5.1.0) objects were generated using methanol-fixed samples from published single-cell data (Hoang, Wang et al. 2020, Hao, Stuart et al. 2024). After normalization and scaling in Seurat, the plot1cell package (v. 0.0.0.9000) was used to visualize the expression of genes of interest across time points and cell clusters (Wu, Gonzalez Villalobos et al. 2022). To visualize clusters with high *npm1a* expression, we queried the dataset in ProteinPaint (v. 2.81.2-2, https://proteinpaint.stjude.org/F/2019.retina.scRNA.html) to generate tSNE plots.

### Pathway Analysis

Gene ontology pathway analysis for molecular function and biological process was conducted using the clusterProfiler package (v. 4.12) on all SASP factors and senescence markers detected from the database in **Table S2** that had a log_2_FC enrichment > 1 and adjusted *p*-value < 0.05 for individual time points. GO terms were subject to a *q*-value cutoff of 0.05 and *p*-values were adjusted using the Benjamini-Hochberg procedure in clusterProfiler. These results were visualized using enrichPlot (v. 1.24) and ComplexHeatmap (v 2.22.0) (Yu, Li et al. 2010, Yu, Wang et al. 2012) and are in **Table S4**.

We used InteractiVenn to compare SASP factor and senescence marker enrichment across multiple datasets (**Table S5**) and generate a “Regeneration-associated Senescence Signature” (RASS) (Heberle, Meirelles et al. 2015). Using a list of SASP factors shared across the four acute damage datasets with multiple collection points (upregulated genes from the GFP^+^ and GFP^-^ conditions for Hoang et al. 2020 were aggregated together) (Hoang, Wang et al. 2020), the STRING database (v. 12.0) was then used to generate a network visualization of potential gene interactions and analyze enriched Reactome pathways (Szklarczyk, Kirsch et al. 2023). Reactome terms were reported if their false discovery rate was ≤ 0.05, minimum signal and signal strength were ≥ 0.01, and had a gene count ≥ 2. Terms were combined if their similarity was ≥ 0.9.

### Animal Care and Husbandry

A mixture of male and female wild-type AB zebrafish was used for experimentation and maintained at 28.0°C on a 12:12hr light-dark cycle. All experiments were performed in accordance with Vanderbilt University Institutional Animal Care and Use Committee (IACUC) approval #M1800200.

### *npm1a* knockdown and NMDA damage

Antisense oligonucleotides (ASOs) were designed to target *npm1a* and modified with 2ʹ-deoxyribonucleotides at central nucleotides flanked on both 5ʹ and 3ʹ sides by 2ʹ-methoxyethyl (MOE)-modified nucleotides (Maimon, Chillon-Marinas et al. 2021)

Additionally, internucleotide linkages were phosphorothioates (PSs) interspersed with phosphodiesters, and all cytosine residues were 5ʹ-methylcytosines. Target sequences for ASOs are listed in **Table 2**. Adult zebrafish aged 5-10 months were anesthetized using a 0.016% Tricaine solution (MESAB, Acros Organics). The sclera of the left eye was cut using a sapphire blade, and 0.5 µL of a 10 mM NMDA solution were intravitreally injected. An equal volume of 1X PBS was used as an injection control for the NMDA damage. *npm1a* ASOs were co-injected with either NMDA or PBS at a concentration of 100 µM and volume of 0.5 µL, with 0.5 µL of 100 µM non-targeting random ASOs serving as controls.

**Table 2:**
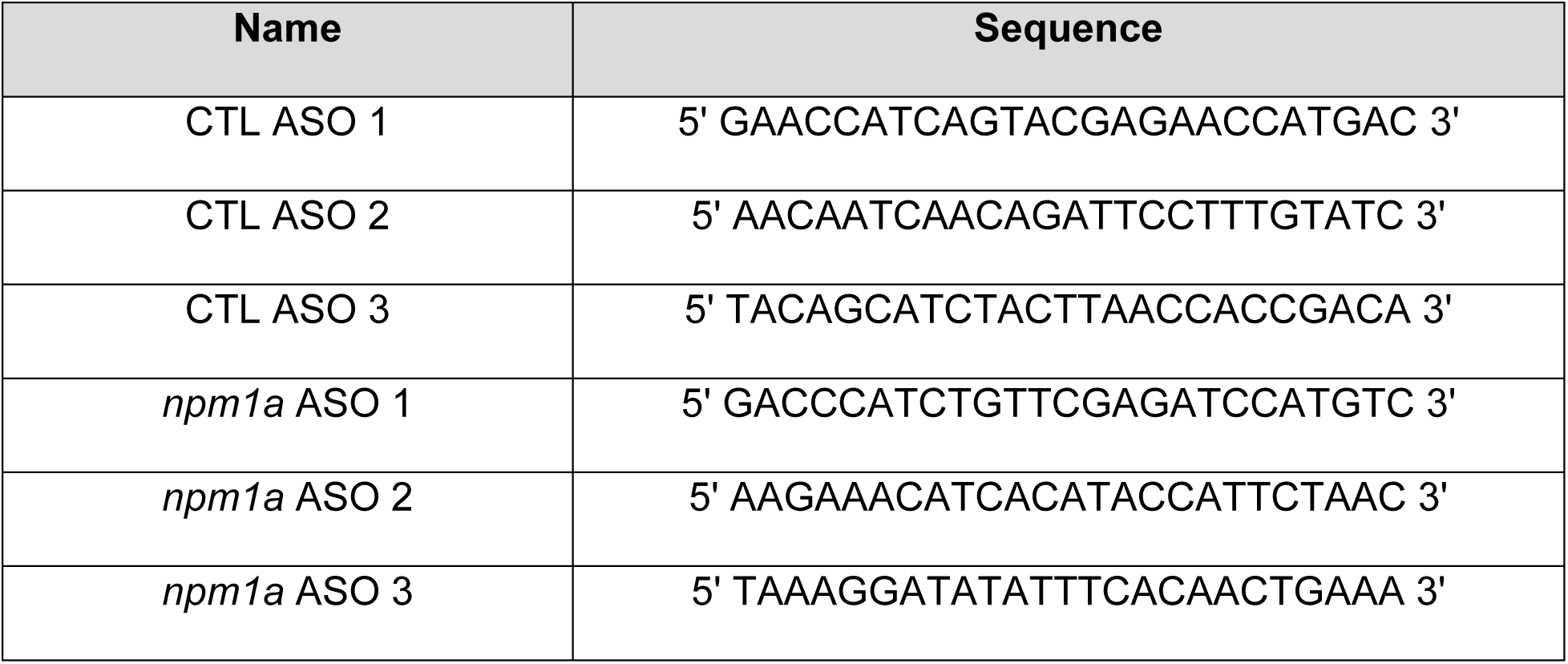
Antisense Oligonucleotide Sequences.

### 4c4 and EdU Staining

Fluorescent staining was performed as previously described (Konar et al 2024). Briefly, eyes were collected and fixed in a 4% paraformaldehyde solution overnight. Eyes were then incubated in a 5% sucrose solution 4 times and kept overnight in 30% sucrose. They were then incubated in a 2:1 solution of Cryomatrix with 30% sucrose solution for 2 hours and embedded in Cryomatrix (Fisher Scientific). Retina sections were generated using a Leica Cryostat to a thickness of 15 µm and collected onto positively charged microscope slides (VWR International). Slides were briefly dried on a heating block and then stored at −80°C. For IHC, slides were warmed on a heating block and then rehydrated in 1X PBS for 30 minutes. Slides were then subjected to antigen retrieval before being allowed to cool to room temperature (RT) (Konar, Flickinger et al. 2024). Blocking solution (3% donkey serum, 0.1% Triton X-100 in 1X PBS) was applied to the slides and incubated for 2 hours at RT. Slides were then stained with 1:500 4c4 antibody (gift from Hitchcock Lab, Univ. of Michigan) overnight at 4°C. After primary antibody incubation, slides were stained with a donkey anti-mouse cy3 antibody (Jackson Labs) at 1:500 in 1% donkey serum, 0.05% Tween-20 in PBS for 2 hours at RT. Slides were then dried and mounted using VectaShield Antifade Mounting Medium with DAPI (Vector Labs).

For EdU staining, zebrafish received an intravitreal injection of 0.5 µL of a 20mM EdU solution at 24 hours post injury ( hpi). Staining was performed using a Click-iT™ Plus EdU Cell Proliferation Kit (555 fluorophore, ThermoFisher, USA) with DAPI nuclear stain.

### Senescence associated β-galactosidase staining

Senescence associated β-galactosidase (SA-βGal) staining was performed on 15µm thick retina tissue sections using a kit from Cell Signaling Technology. Sections were rehydrated in PBS and a SA-βGal staining solution at pH 5.9-6.1 was administered overnight at 37°C. Samples were then washed with PBS and treated with a nuclear staining solution containing 1:1000 TO-PRO-3 (ThermoFisher) in 1X PBS. Slides were then dried and mounted using VectaShield AntiFade Mounting Medium (Vector Labs) and sealed with nail polish.

### Quantitative reverse transcription polymerase chain reaction (qRT/PCR)

qRT/PCR was performed on retinas dissected from NMDA treated eyes as previously described (Kent, Kara et al. 2021, Konar, Flickinger et al. 2024). Briefly, retinas were collected into Trizol (Invitrogen), and mRNA was converted to cDNA using an Accuscript High-Fidelity 1^st^ Strand cDNA Synthesis Kit. qRT/PCR of the cDNA samples was performed using SYBR Green (Bio-Rad Laboratories). Samples were run on a 96-well plate using a Bio-Rad CFX96 Real-time System. qPCR runs were normalized to 18S rRNA levels and analyzed using the ΔΔCt method. The primer sets used are listed in **Table 3**.

**Table 3.**
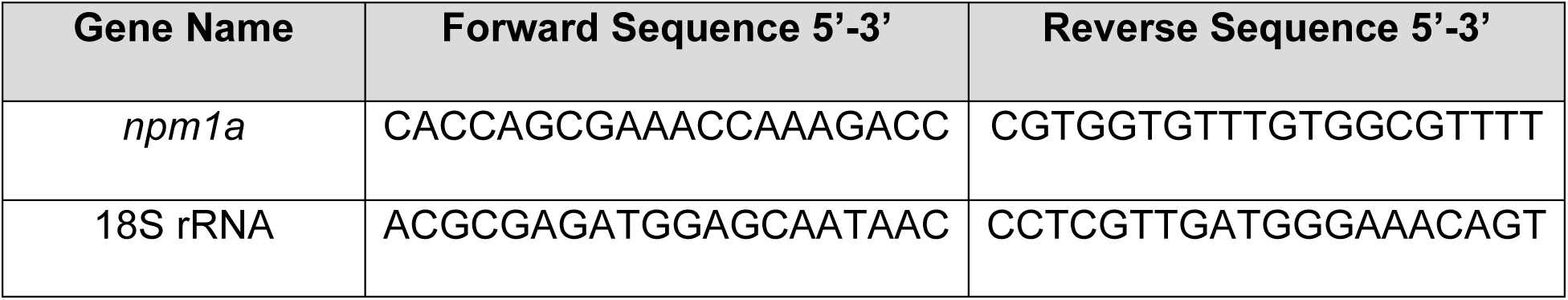
Primer sequences used for qRT/PCR of *npm1a* ASO treated retinas.

### Imaging and Image Analysis

Slides stained with SA-βGal and the nuclear stain TO-PRO-3 were imaged using a Nikon AZ100M Widefield microscope in the Vanderbilt Imaging Core (CISR). AZ100M images were first processed using NIS-Elements Viewer 5.21 and then subsequently analyzed using ImageJ. Slides stained with IHC or EdU and DAPI were imaged using a Zeiss 880 Confocal Microscope through the Vanderbilt CISR Core. Zeiss 880 images were processed using Zeiss Blue software and subsequently analyzed using ImageJ.

Experiments involving cell number quantification were evaluated in an unbiased and blinded manner. Two retina sections containing both dorsal and ventral regions of the retina and flanking the optic nerve region by 200 µm on each side were counted and averaged from each injected eye. Significance was calculated using a two-way ANOVA with Tukey’s post-hoc test for inter-group comparisons in GraphPad Prism 10.4.0. All data represented are reported as the average ± standard error of the mean (s.e.m). The sample size for each experiment is stated in the respective figure legends.

## Results

### Dynamic SASP factor expression after acute damage in the zebrafish retina

NMDA and constant intense light damage are two common paradigms used to acutely damage the retina and trigger a MG-derived regeneration response (Vihtelic, Soverly et al. 2006, Karl, Hayes et al. 2008). To identify differentially expressed SASP factors during retina regeneration, we analyzed published and in-house datasets (Kramer, Gurdziel et al. 2021) to visualize trends in SASP factor expression during and after acute damage to the retina. The light damage dataset utilized an acute light damage paradigm featuring a 5 day dark adaptation followed by phototoxic lesion formation and 3 days of halogen light exposure, leading to robust degeneration of photoreceptors and activation of MG (Kramer, Gurdziel et al. 2021). Our in-house NMDA dataset utilized intravitreal injections of NMDA to induce retinal ganglion cell death and activation of MG (Karl, Hayes et al. 2008). Both data sets featured collections at or before 3 days post injury (dpi), around 10-12 dpi, and then near the completion of regeneration at 20-28 dpi.

During light damage, we detected an almost immediate upregulation of SASP factors between 24-36 hpi, most notably thioredoxin b (*txnb*), annexin (*anxa2a*), nucleophosmin 1a (*npm1a*), and matrix metallopeptidase 9 (*mmp9*) (**Figure 1A-B**). By 72 hpi, some of the initially upregulated SASP factors began to abate, including *npm1a* and the tubulins *tubb6*, *tuba1b*, and *tuba5* (**Figure 1C,H)**. A more prominent shift in SASP factor expression was observed at 5 dpi, with most SASP factors starting to return to baseline (with the exception of complement factors *c3a.4* and *c3a.1* and aldo-keto reductase *akr1a1a*), a trajectory which continued through 28 dpi (**Figure 1D-H)**. This decrease in SASP factor expression after initial enrichment highlights the dynamic and transient gene expression patterns that occur after damage.

**Figure 1.**
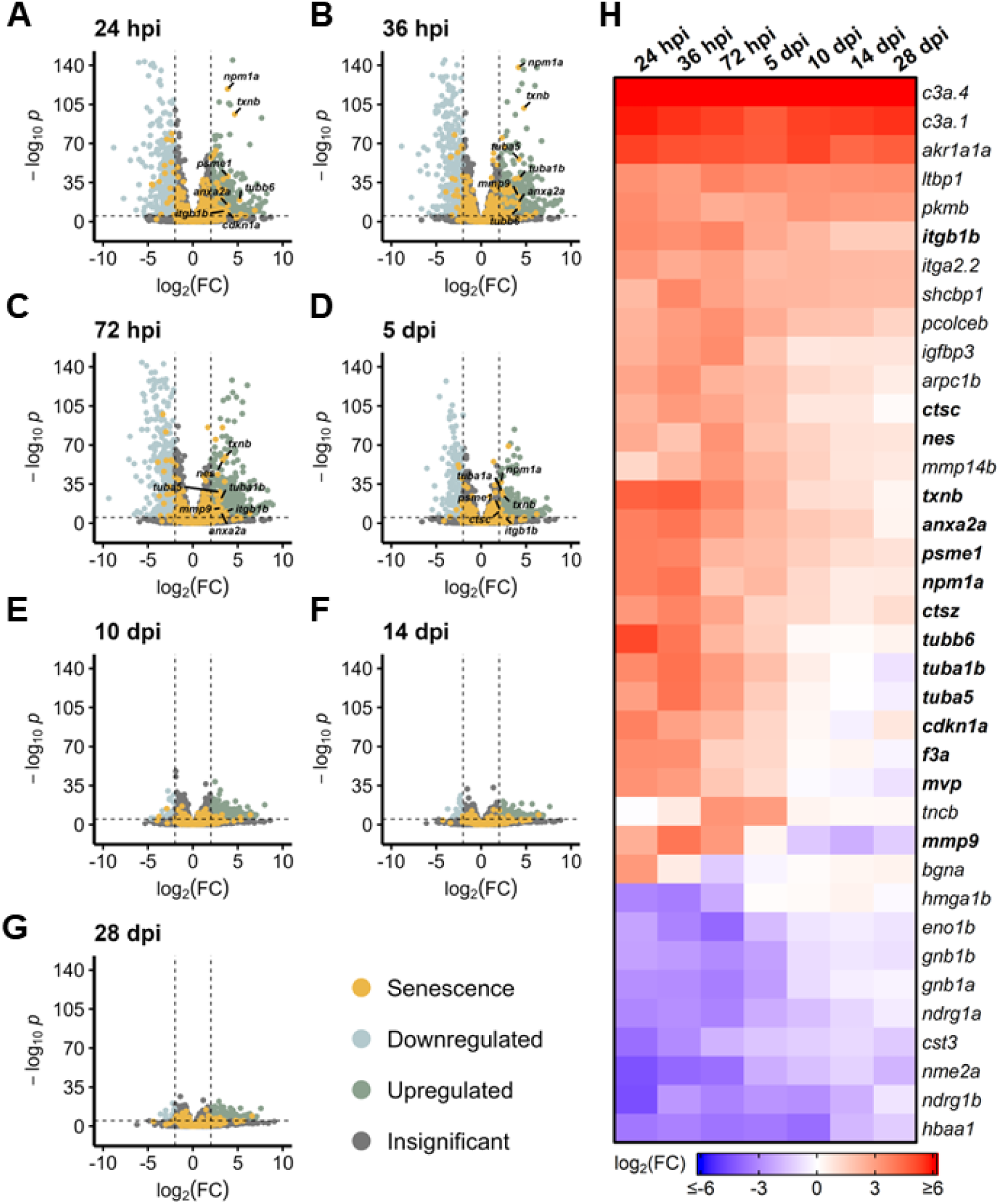
Differentially expressed SASP factors after acute light damage. Bulk RNAseq was performed on whole zebrafish retinas after acute light damage and compared to undamaged controls. (A-G) Volcano plots depicting up- and downregulated SASP factors and senescence markers at the indicated times post damage. Dashed lines represent values with a |log_2_FC| > 2 and *p*-values < 10^-6^. Top significantly upregulated factors are labeled for each time point. (H) Heatmap showing enrichment of SASP factors at the indicated times post light damage. SASP factors and senescence markers were included in the heatmap if they had a log_2_FC > 3 with *p*-values < 10^-6^ for at least one time point; bolded genes are part of the Regeneration-associated Senescence Signature (RASS).

After NMDA damage, we also observed an early burst of differentially expressed SASP factors at 3 dpi, led by *mmp9*, *anxa2a*, and *txnb* (**Figure 2A**). Other SASP factors that were elevated early after NMDA damage include the cell cycle gene tubulin alpha 5 (*tuba5*) and the transcription factor *jun*. At 12 dpi, we observed a decrease in SASP factor expression that continued through 20 dpi (with the exception of *anxa2a*), similar to the trend observed after acute light damage (**Figure 2B-D**). Taken together, both light and NMDA damage show conserved differential gene expression patterns, with several common SASP factors showing up-regulation during the early stages of regeneration.

**Figure 2.**
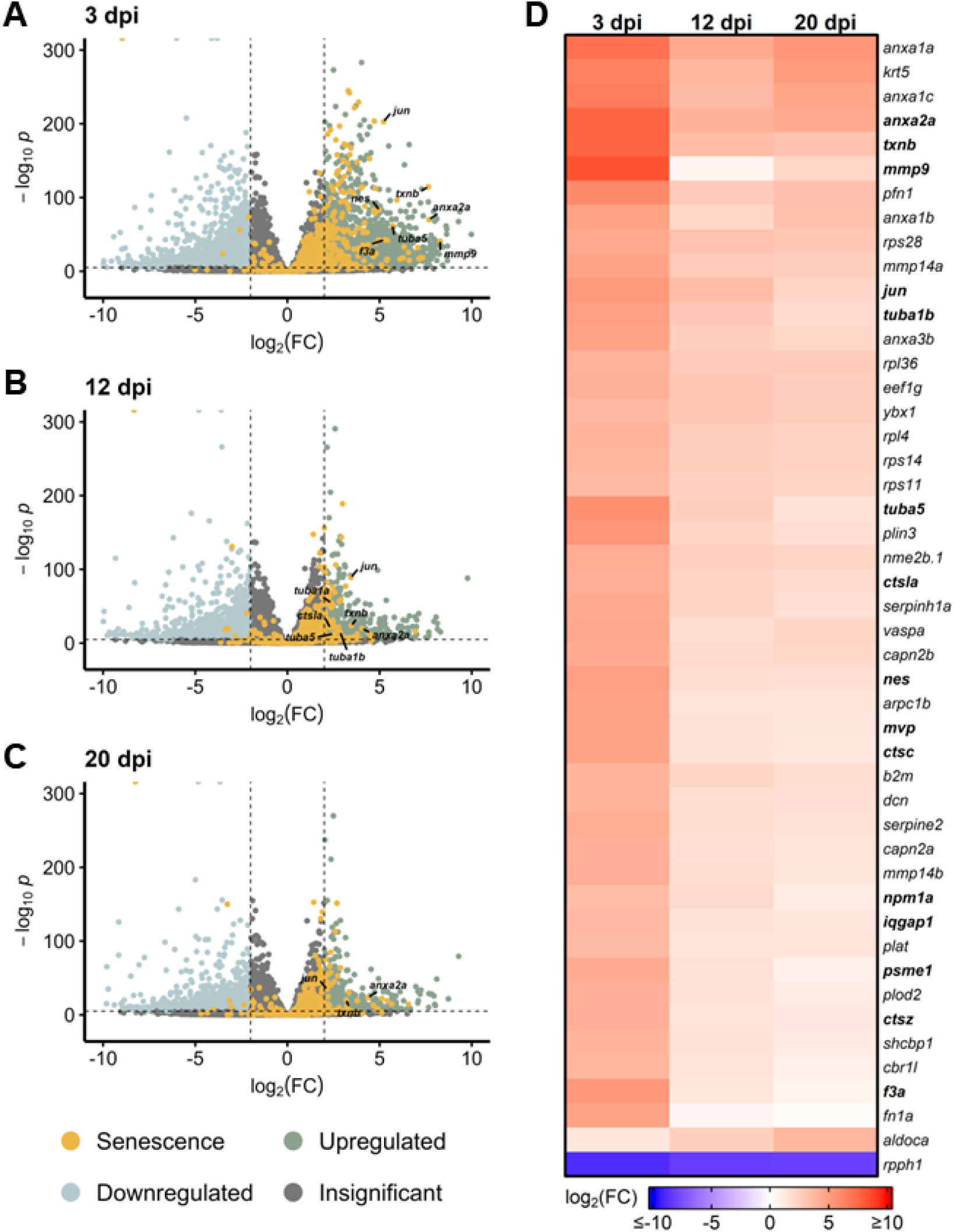
Differentially expressed SASP factors after NMDA damage. Bulk RNAseq was performed on whole zebrafish retinas after intravitreal injections of NMDA and compared to undamaged controls. (A-C) Volcano plots showing differentially expressed SASP factors at the indicated times post damage. Dashed lines represent values with a |log_2_FC| > 2 and *p*-values < 10^-6^. Top significantly up-regulated factors are labeled for each time point. (D) Heatmap showing enrichment of SASP factors at the indicated times post NMDA damage. SASP factors were included in the heatmap if they had a log_2_FC > 3.5 with *p*-values < 10^-20^ for at least one time point; bolded genes are part of the RASS.

### Gene ontology of SASP factors after acute damage shows variable enrichment of gene function and process over time

SASP factor expression is thought to directly influence the microenvironment. After acute light damage, Gene Ontology (GO) analyses showed an enrichment of SASP factors implicated in the regulation of endopeptidase activity and the binding of cell adhesion molecules (**Figure 3A)**. SASP factors with molecular functions related to collagen synthesis were also upregulated before 5 dpi with additional functions related to ECM remodeling and binding enriched throughout regeneration. Changes in these ECM-related functions are further supported by biological process enrichment analysis (**Figure S2, Table S4**). Similar ECM-related processes were enriched at 3 days post NMDA damage (**Figure S3**), though molecular function enrichment was more heterogeneous, with elevated protein folding functions across the time course (**Figure 3B**). The changing enrichment of specific GO functions and processes carried out by SASP factors between early and late regeneration suggest that dynamic changes in the SASP secretome during regeneration modulate the local microenvironment to facilitate regeneration.

**Figure 3.**
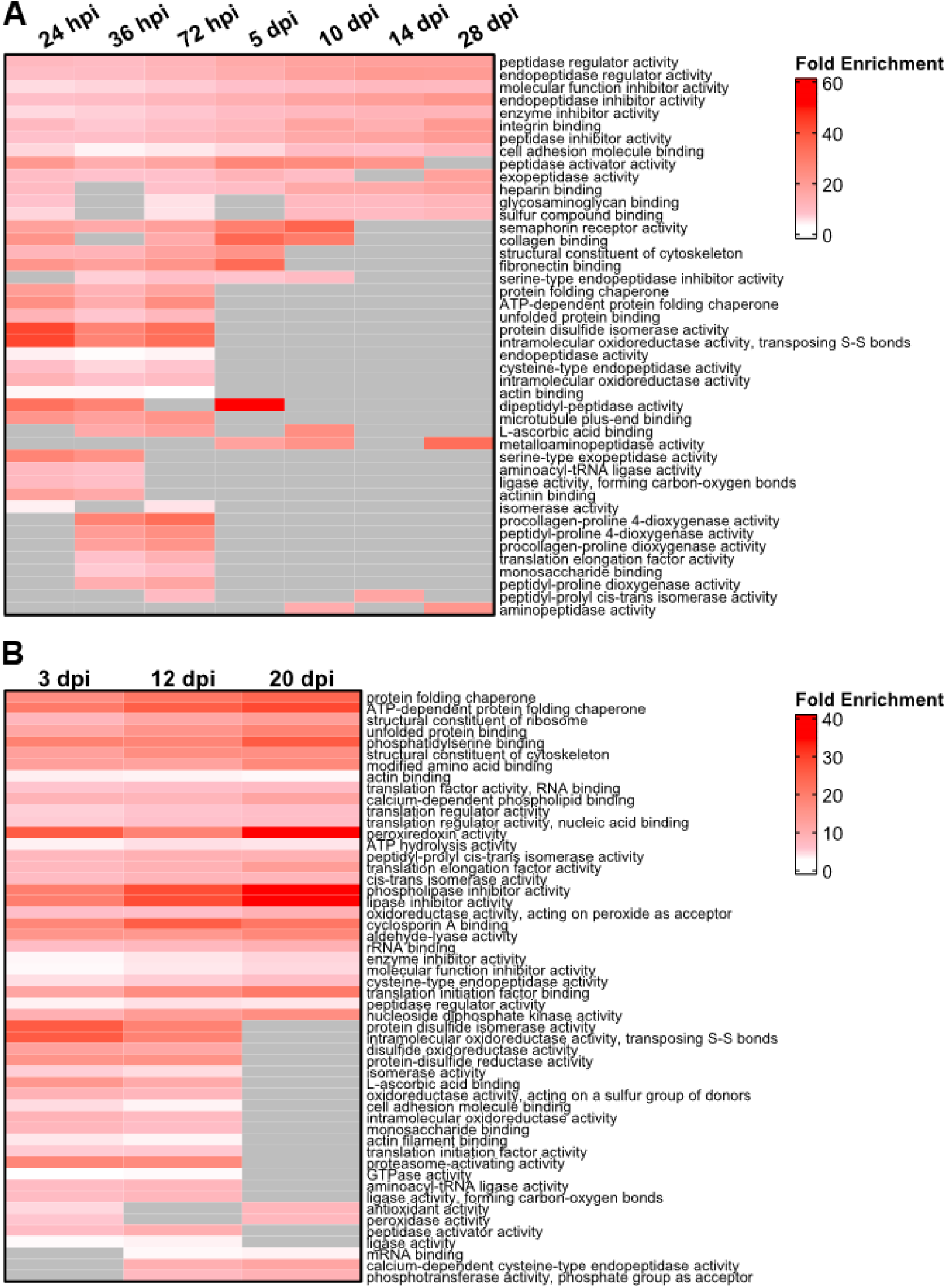
Gene Ontology (GO) analysis of differentially expressed SASP factors after light and NMDA damage. (A) GO analysis of the molecular function of the most enriched SASP factors in light damaged retinas showed enrichment for genes involved in the binding of integrins and fibronectin, as well as isomerase, peptidase, and chaperone function. (B) GO analysis of the molecular function of SASP factors in NMDA damaged retinas showed enrichment of genes responsible for lipase inhibitor activity, proteasome activation, and protein chaperone activity. SASP factors were included in the overrepresentation analysis if they had an adjusted p-value < 0.05 and log_2_FC > 1. GO terms were included in each heatmap if they were enriched for at least two time points. Gray cells indicate time points at which the respective term was not enriched.

### Conserved SASP factor expression in Muller glia after acute damage

To determine whether there is a MG-specific SASP profile, we examined bulk RNAseq data sets from sorted MG in *Tg[gfap:eGFP]* and *Tg[gfap:GFP]* zebrafish that express GFP in MG (Bernardos and Raymond 2006). Transcriptome studies were performed on MG during early stage (4-36 hpi) regeneration after both light and NMDA damage (Hoang, Wang et al. 2020). We detected a robust SASP profile after both damage modalities compared to undamaged controls, with increased expression of several SASP factors that were also observed in Figures 1 and 2 (*anx2a, mmp9, txnb, tuba1b, tubb6*) (**Figure 4A-H**). When directly comparing light and NMDA damage in MG at early stages, the SASP profile showed high levels of conservation of factors and this pattern held when comparing GFP^+^ MG to GFP^-^ cells (**Figure 4I**, **Figure S4**). This indicates that the SASP profile in MG shows similar conservation to whole retina transcriptomic analysis of SASP factors after damage, regardless of the acute damage type.

**Figure 4.**
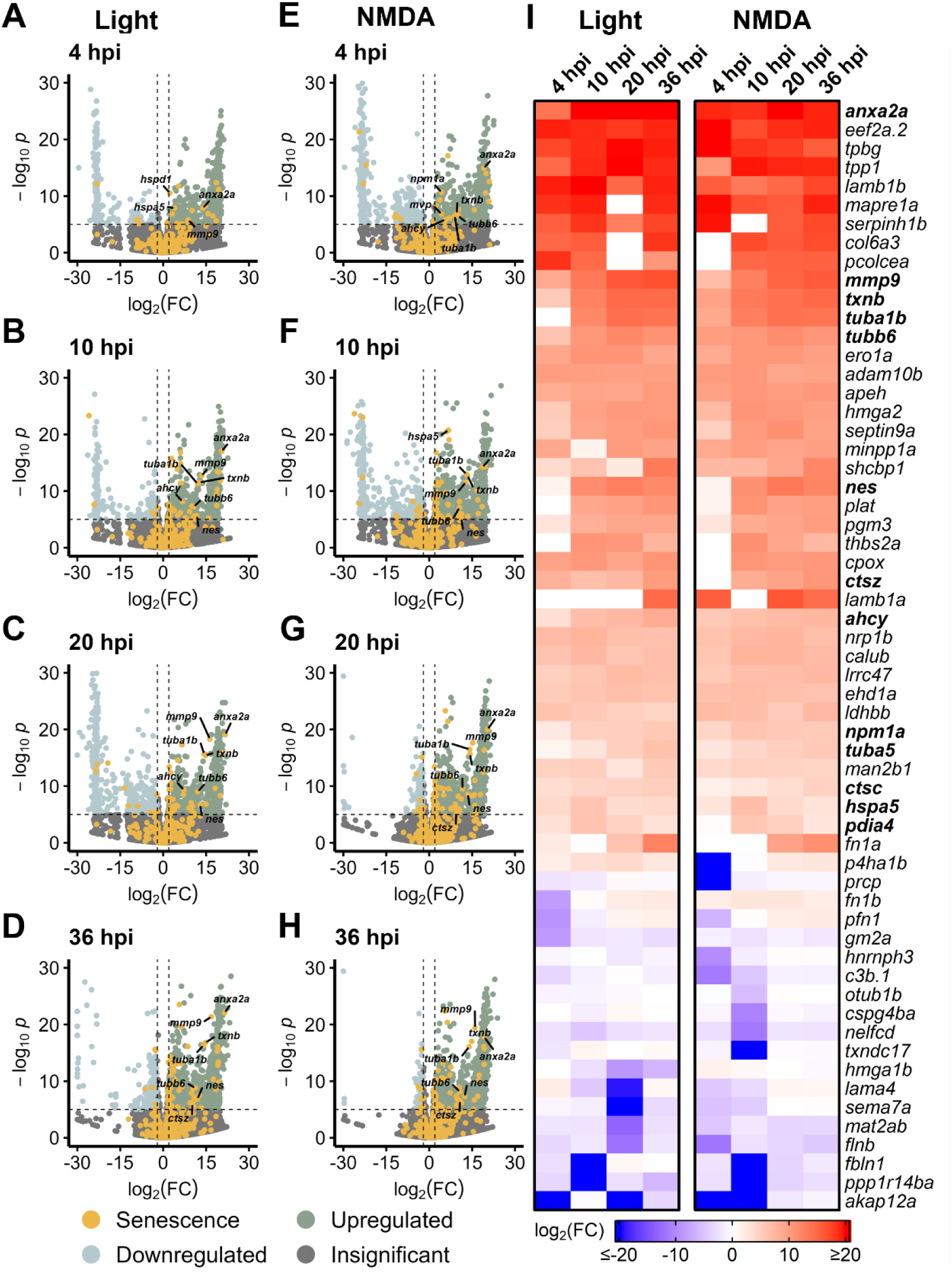
Upregulation of SASP factors in Müller glia after light and NMDA damage. Bulk RNAseq was performed on *gfap*^+^ Müller glia after either NMDA or light damage compared to undamaged controls. (A-D) Volcano plots showing differentially expressed SASP factors at the indicated times post light damage. (E-H) Volcano plots showing differentially expressed SASP factors at the indicated times post NMDA damage. Dashed lines represent values with a |log_2_FC| > 2 and *p*-values < 10^-6^. Top significantly up-regulated factors are labeled for each time point. (I) Heatmap showing enrichment of SASP factors at the indicated times post light or NMDA damage. SASP factors were included in the heatmap if they had a log_2_FC > 5 with *p*-values < 10^-6^ for at least one time point; bolded genes are part of the RASS.

### Enrichment of distinct SASP factors in a chronic model of retina damage

Chronic damage mutants have shown drastically different regeneration patterns and gene expression compared to acute models, prompting the question of whether SASP factor expression patterns vary between acute and chronic damage (Gorsuch and Hyde 2013, White, Sengupta et al. 2017, Fogerty, Song et al. 2022, Iribarne and Hyde 2022, Grabinski, Parsana et al. 2023, Kramer, Carthage et al. 2023). Two models of chronic damage use ciliopathy mutants (*bbs2^-/-^* and *cep290^-/-^*) that impact photoreceptors, with *bbs2^-/-^* mutants losing photoreceptors by 4 months post fertilization (mpf) and *cep290^-/-^* mutants losing photoreceptors by 6 mpf (Grabinski, Parsana et al. 2023). We analyzed gene expression patterns (Grabinski, Parsana et al. 2023) in these mutants and found that *bbs2^-/-^* mutants showed a much more robust differential expression profile of SASP factors than *cep290^-/-^* mutants. Differentially expressed SASP factors in the *bbs2^-/-^* mutant were similar to factors identified after acute damage, including *mmp9*, *anxa2a*, *txnb*, and *tuba5* (**Figure 5A**). Both *bbs2^-/-^ and cep290^-/-^* mutants showed increased expression of the senescence marker *jun* and the SASP factor *txnb* (**Figure 5B**). Overall, *bbs2^-/-^* mutants showed more pronounced elevation of almost every SASP factor than the *cep290^-/-^* mutants, except for the inflammatory cytokine IL8 (*cxcl8a)* and the transcription activator *stat1b* (**Figure 5C**). GO analysis of both mutants showed enrichment for SASP factors regulating endo- and exo-peptidase activity in *bbs2^-/-^* mutants. In *cep290^-/-^*, there was greater enrichment of factors influencing protein kinases and the GO term enzyme inhibition (**Figure S5**). Given the variation in the damage response between chronic and acute models of retinal damage, the similarities between enriched SASP factors in the *bbs2^-/-^* mutants compared to light and NMDA acute damage models support conserved microenvironmental SASP cues between both chronic and acute damage models.

**Figure 5.**
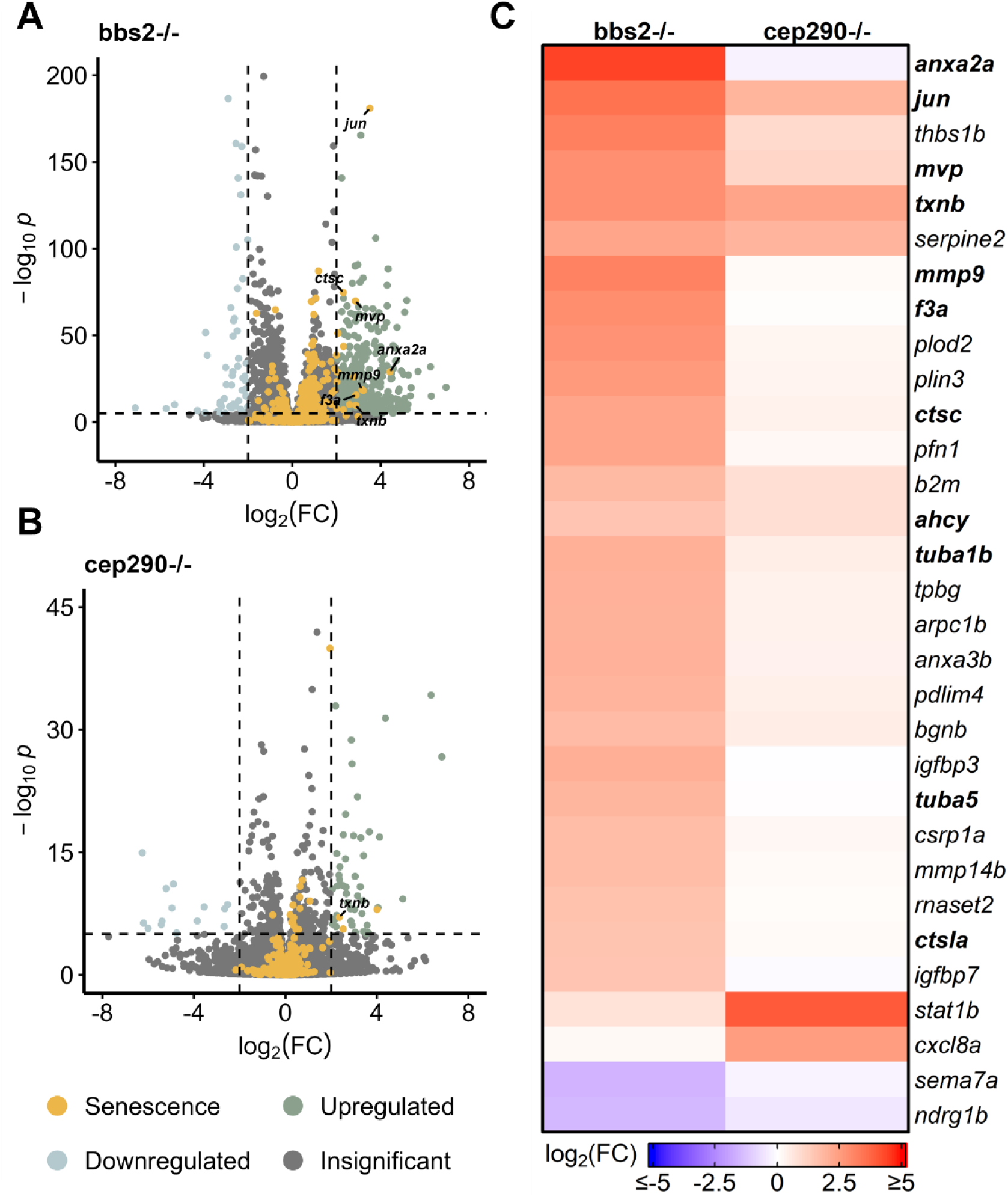
Differentially expressed SASP factors in genetic mutants with chronic photoreceptor damage. Bulk RNAseq was performed on whole zebrafish retinas from *bbs2^-/-^* and *cep290^-/-^* ciliopathy mutants. (A) Volcano plots of differentially expressed SASP factors in *bbs2^-/-^ or cep290^-/-^* mutants at 6 mpf. Dashed lines represent values with a |log_2_FC| > 2 and *p*-values < 10^-6^. Top significantly up-regulated factors are labeled for each time point. (B) Dot plot analysis of SASP factors with |log_2_FC| > 1 shows enrichment for genes involved in peptidase, enzymatic processes, and lipase activity in *bbs2^-/-^* mutants. *cep290^-/-^* mutants also showed enrichment for genes regulating enzyme inhibition and protein kinase inhibition, but at a far lower enrichment than in *bbs2^-/-^* mutants. (C) Heatmap showing differentially expressed SASP factors with a log_2_FC > 1.5 and *p*-values < 10^-6^ in the indicated mutants; bolded genes are part of the RASS.

To further understand differences between chronic and acute damage models, we evaluated a chronic light damage model of retina damage (Kramer, Carthage et al. 2023). Expression of SASP factors in chronic light damage was upregulated most around 36-72 hpi, a smaller window for elevated SASP expression than in acute light damage (**Figure S6A-G**). Chronic light damage models showed an overall lesser expression of SASP factors compared to acute damage. However, expression of the SASP factors *ctsba, txnb, psme1*, and the senescence marker *jun* were still appreciably upregulated in this damage model (**Figure S6H**). The relatively minor upregulation of SASP factors when comparing chronic to acute light damage may underscore differences in the timing or mechanisms of repair between the two models.

### Regeneration-based SASP signatures after acute and chronic retinal damage

To determine whether there is a regeneration specific signature among SASP factors expressed post-damage, we compared the upregulated SASP factors from both acute and chronic damage models. After acute damage, we identified 29 SASP factors and 2 senescence markers that overlapped between all 4 datasets (Hoang, Wang et al. 2020, Kramer, Gurdziel et al. 2021). We refer to these common factors as the “Regeneration-associated Senescence Signature” (RASS) (**Figure 6A, Table S5**). We then compared that RASS list to upregulated factors from the 3 chronic conditions (Grabinski, Parsana et al. 2023, Kramer, Carthage et al. 2023) and identified 2 SASP factors and 1 senescence marker enriched across all the datasets (**Figure 6B**). These include thioredoxin b (*txnb*), major vault protein (*mvp*), and the jun proto-oncogene (*jun*) (**Table S5**). We then examined the protein-protein association network of the RASS and found that 25 of the 31 factors had conserved or predicted protein-protein relationships between one another, particularly between the tubulins and heat shock proteins (**Figure 6C**). To better understand the biological pathways and reactions shared between the RASS constituents, we performed Reactome pathway analysis (Szklarczyk, Kirsch et al. 2023) and found enrichment in antigen presentation, cell transport, cell cycle regulation, immune system regulation, and aggrephagy—a process that selectively removes protein aggregates via macroautophagy (**Figure 6D, Table S6**) (Lamark and Johansen 2012). This core set of SASP factors represents potential regeneration-specific genes secreted by senescent cells after damage.

**Figure 6.**
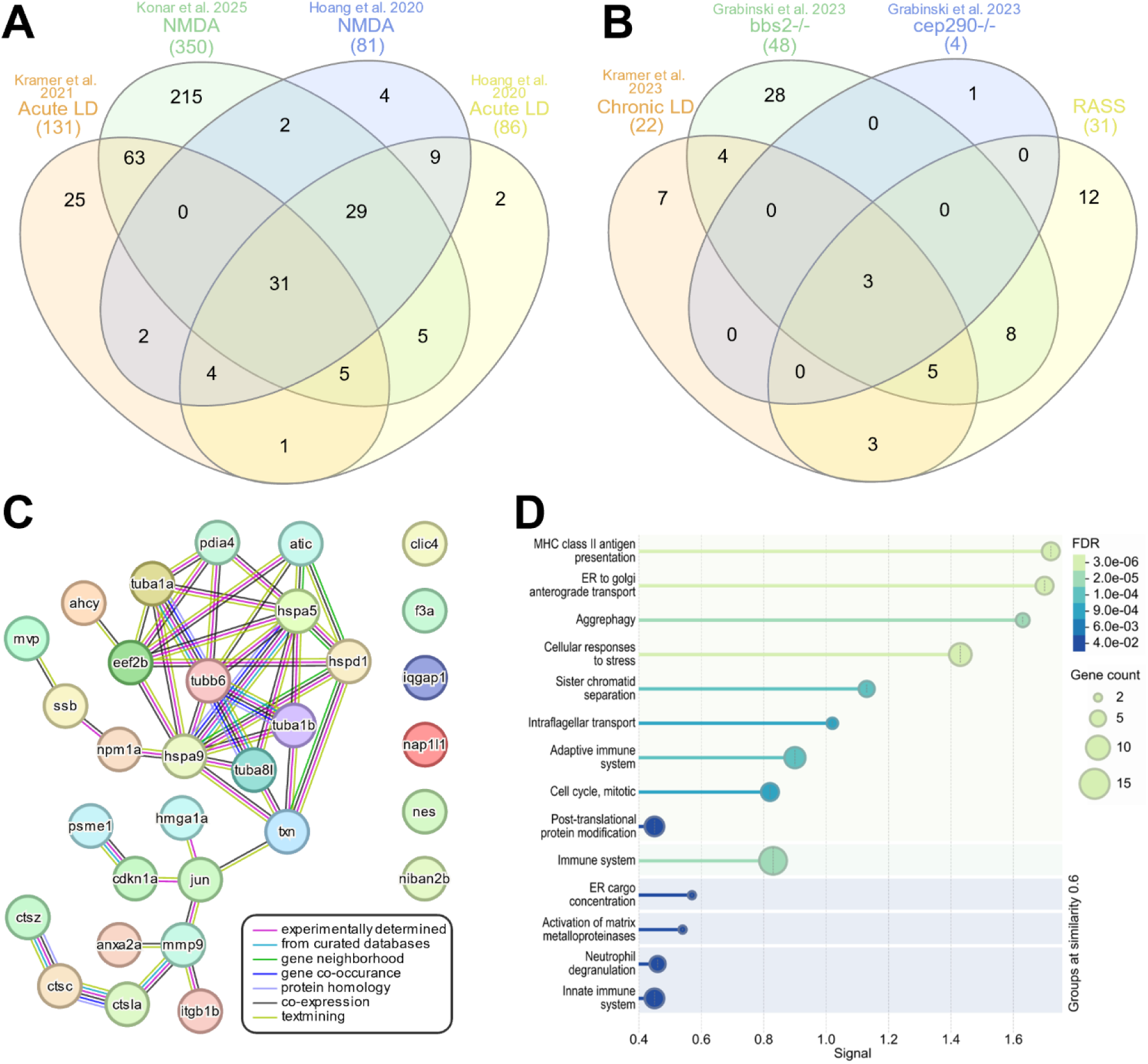
Regeneration specific secretome signature. InteractiVenn based analysis was used to determine the presence of a conserved secretome signature. (A) Analysis of upregulated SASP factors and senescence markers from 4 acute damage datasets identified 31 conserved genes, referred to as the Regeneration-associated Senescence Signature (RASS). (B) Analysis of upregulated SASP factors and senescence markers in chronic damage datasets identified 3 conserved genes. (C) STRING analysis of the RASS showed both predicted and conserved protein-protein relationships between 25 of the 31 genes in the RASS. (D) Reactome analysis of the RASS showed enrichment in processes related to antigen presentation, cellular transport, immune system processing, and aggrephagy.

### SASP factor expression enriched in glial cells

To begin to assess what cell types are expressing SASP factors during regeneration, we decided to focus on 6 key factors. *mmp9*, *npm1a*, and *anxa2a* showed conserved positive enrichment during early-stage regeneration that persisted across damage modalities and datasets. Thioredoxin B (*txnb*) is a redox protein that is involved in cell-cell communication and stress response and is elevated during muscle regeneration (Vezzoli, Castellani et al. 2011). *hmga1a* is a non-histone chromatin protein (Harrer, Lührs et al. 2004) that is upregulated after NMDA and light damage (Hoang, Wang et al. 2020). Lastly, *tuba1a* was chosen because it is a well-known retinal progenitor cell marker and SASP factor involved in MG-derived retina regeneration (Ramachandran, Reifler et al. 2010). We analyzed the cellular localization of these factors based on scRNAseq using Seurat and the plot1cell package (Hoang, Wang et al. 2020, Wu, Gonzalez Villalobos et al. 2022, Hao, Stuart et al. 2024). *mmp9* expression localized to both resting and activated MG, as well as in microglia (**Figure 7A**). *txnb* expression localized primarily to resting and activated MG after both NMDA and light damage, but localization was also observed in cones and RPE cells after light damage (**Figure 7B**). For both *mmp9* and *txnb*, there was an increase in the percentage of cells in each cluster expressing these genes throughout early regeneration.

**Figure 7.**
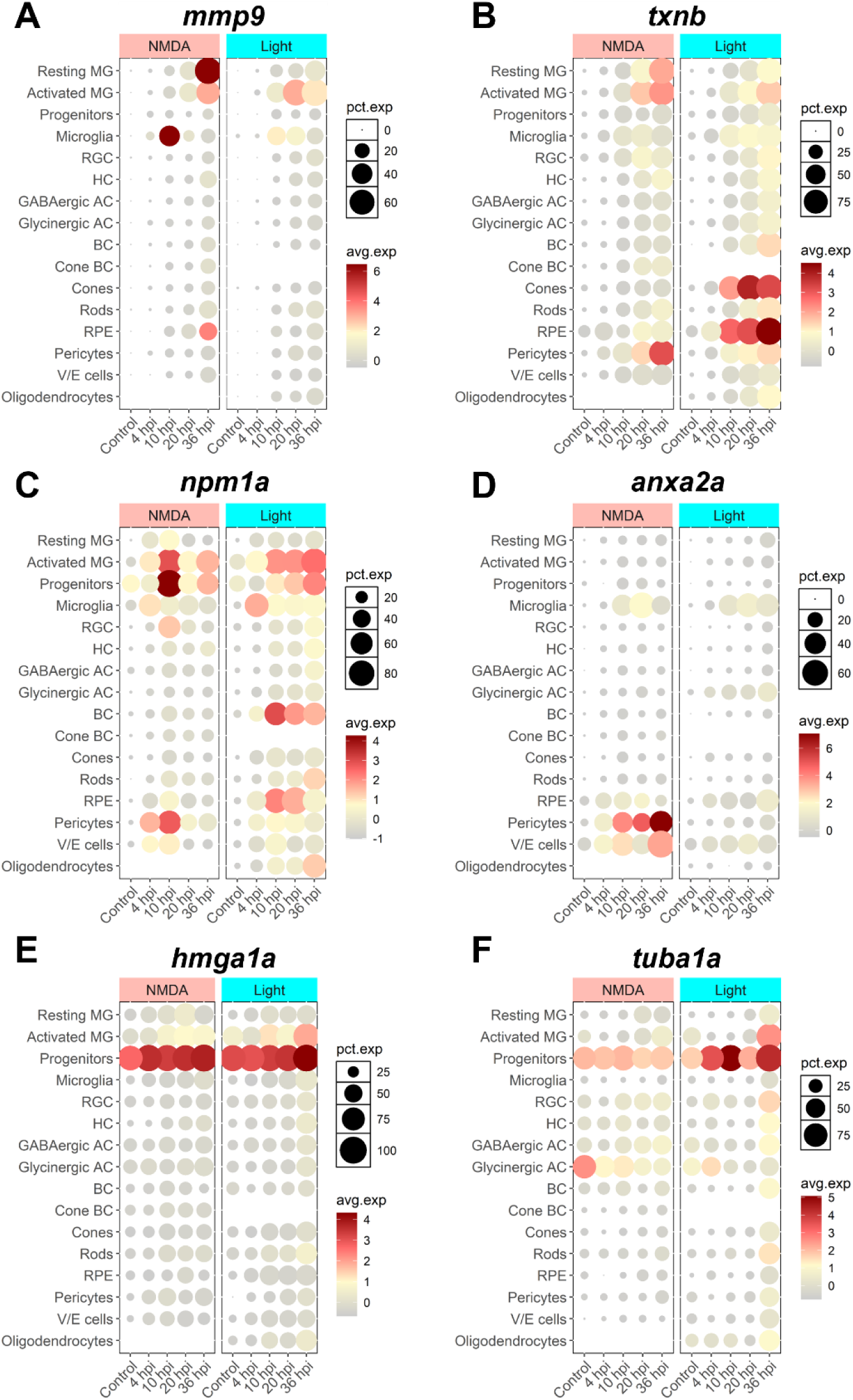
scRNAseq visualization of SASP factors after NMDA and light damage. Localization of SASP factor expression based on scRNAseq data from dissociated retinas was visualized using Seurat and the plot1cell package. (A) *mmp9* expression was elevated in MG, microglia, and RPE. (B) *txnb* expression was elevated in MG and pericytes in both damage models, and cones and RPE in light damage only. (C) *npm1a* expression was elevated in MG, progenitors, microglia, pericytes, and bipolar cells. (D) *anxa2a* expression was elevated in microglia, pericytes, and vascular/endothelial cells. (E-F) *hmga1a* and *tuba1a* expression were largely elevated in progenitor cells. The average normalized expression within each cluster and the percentage of cells in each cluster expressing each gene of interest is shown. MG = Müller glia, RGC = retinal ganglion cell, HC = horizontal cell, AC = amacrine cell, BC = bipolar cell, RPE = retinal pigment epithelium, V/E cells = vascular endothelial cells.

Expression of *npm1a* localized to activated MG and progenitor cells, microglia, pericytes, with limited expression detected in bipolar cells (**Figure 7C**). *anxa2a* expression localized primarily to microglia, pericytes, and vascular-endothelial cells (**Figure 7D**). Finally, *hmga1a* and *tuba1a* expression localized almost exclusively to retinal progenitor cells (**Figure 7E**). Overall, the presence of SASP factors in both activated MG and microglia suggests potential crosstalk between glial cell types after damage. This supports a model whereby senescent cells might exert a paracrine influence on MG reactivity.

### Knockdown of Nucleophosmin 1a (*npm1a*) inhibits retina regeneration

Among SASP factors, *npm1a* was significantly upregulated in activated MG and progenitor cells (**Figure 8A**). To assess the role of *npm1a* in retina regeneration, we designed antisense oligonucleotides (ASOs) that target *npm1a* and intravitreally injected them into the retina of adult *Tg[tuba1a:eGFP]* zebrafish in the presence or absence of NMDA damage. We confirmed that the ASOs reduced the expression of *npm1a* via qRT/PCR (**Figure 8B**) and then examined the effects of *npm1a* depletion on senescence and retina regeneration. We found that *npm1a* knockdown decreased the number of senescence associated β-galactosidase^+^ senescent cells after NMDA damage (**Figure 8C**). We also observed a decrease in the number of 4c4^+^ immune cells after *npm1a* knockdown in the damaged retina (**Figure 8D**). Finally, we found that MG-derived proliferation after NMDA damage was inhibited by knockdown of *npm1a* (**Figure 8E-F**). During regeneration, PCNA+ proliferating cells typically form clusters associated with MG in the Inner Nuclear Layer (INL) (Rajaram, Harding et al. 2014). When we counted the number of clusters after NMDA damage, we found that knockdown of *npm1a* led to a significant decrease in PCNA+ cells in the INL, indicating impaired MG reactivity in the absence of *npm1a* (**Figure 8E-F**). Taken together, these data show that depletion of *npm1a* inhibits regeneration-associated proliferation after acute damage, suggesting that it is a pro-regenerative factor that is part of the SASP response influencing MG-derived regeneration.

**Figure 8.**
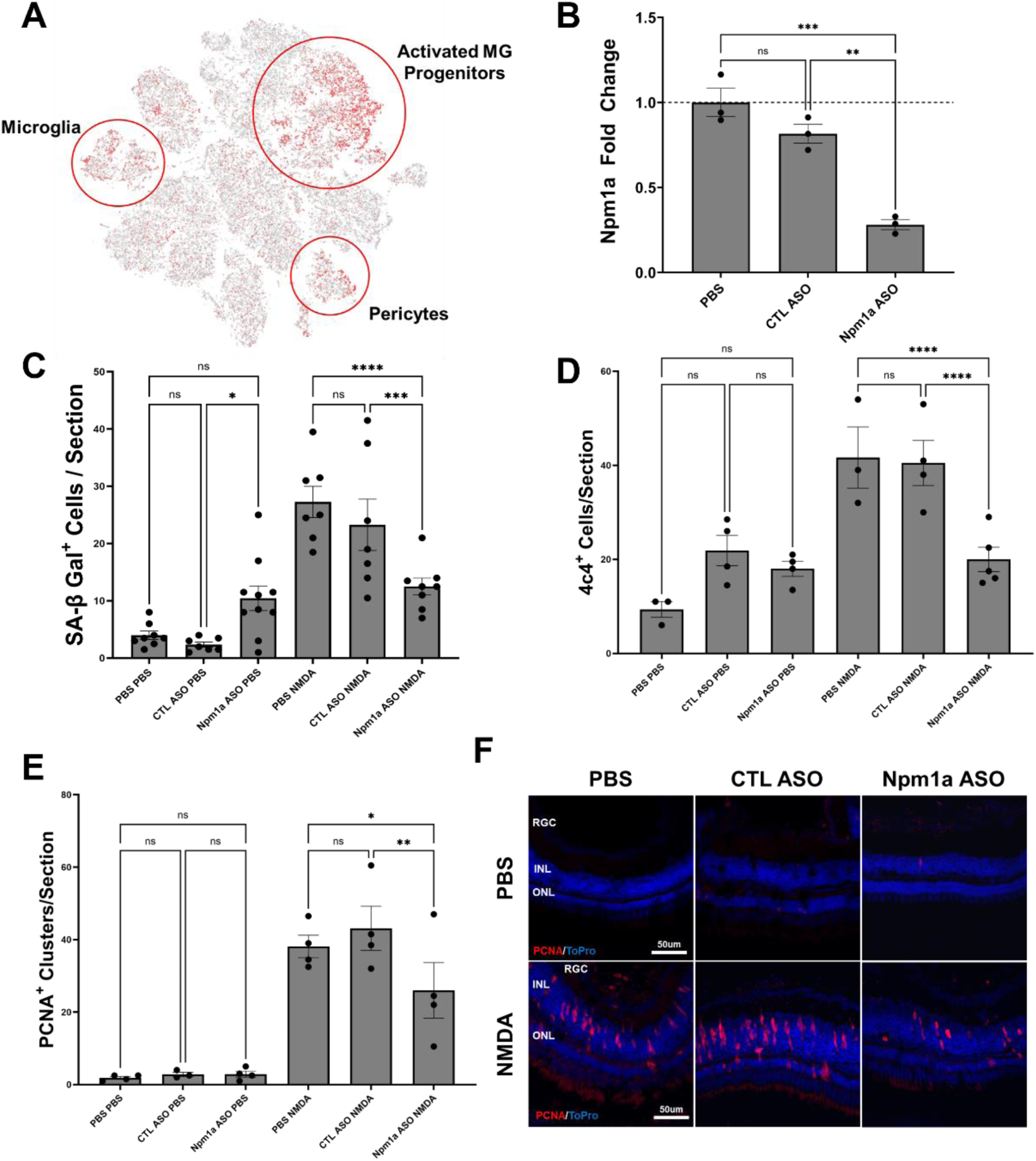
Depletion of Nucleophosmin 1a (*npm1a*) inhibits senescent cell accumulation and proliferation after NMDA damage. (A) Localization of *npm1a* expression by scRNAseq analysis after NMDA damage. (B) Depletion of *npm1a* by intravitreal injection of antisense oligonucleotides (ASOs). Control retinas were injected with either PBS or a mismatch control ASOs, and knockdown of *npm1a* was confirmed using qRT/PCR. (C) Depletion of *npm1a* inhibited accumulation and detection of senescent cells (SA-βgal^+^) at 3 dpi after NMDA damage. (D) Depletion of *npm1a* reduced detection of 4c4^+^ immune cells at 3 dpi after NMDA damage. (E-F) Depletion of *npm1a* inhibited proliferation after NMDA damage at 3 dpi as measured by PCNA immunostaining and counting of MG-associated clusters in the Inner Nuclear Layer (INL) (n=3-6, * = p < 0.05, ** = p < 0.01, *** = p < 0.001, **** = p < 0.0001).

## Discussion

Here, we analyzed transcriptomic data to identify differentially expressed SASP factors using acute and chronic retina damage models in zebrafish. Among these, *mmp9, npm1a, txnb, hmga1a*, and *anxa2a* were the most upregulated SASP factors common to acute and chronic damage models. Of the SASP factors that were the most often upregulated, *mmp9* has been shown to influence retina, nerve, bone, muscle, and limb regeneration by modulating the inflammatory state and influencing ECM degradation (Yang, Gardiner et al. 1999, Kim, Remacle et al. 2012, Wang, Yu et al. 2013, Silva, Nagashima et al. 2020). *hmga1a* is a non-histone DNA binding protein that can modulate chromatin and cellular stemness via induction of pluripotency factors such as *sox2*, *oct4*, *cmyc,* and *lin28* (Parisi, Piscitelli et al. 2020). It has recently been shown to interact with MG after damage, supporting a role as a pro-regenerative factor (Hoang, Wang et al. 2020). Lastly, nucleophosmin 1a (*npm1a*) is a multi-functional protein with roles in genome stability, ribosome biogenesis, cell cycle regulation, and apoptosis (Okuwaki 2008, Lindström 2011). It is often used as a diagnostic factor for acute myeloid leukemia, but its role in the retina is mostly unknown except for its interaction with *chx10*/*vsx2* in MG (Ouchi, Baba et al. 2012). We found that depletion of *npm1a* inhibits MG-derived regeneration. *npm1a* is expressed in activated MG, progenitor cells, microglia, and pericytes, supporting roles in regeneration, modulation of inflammation, and regulation of the blood-retinal barrier (BRB). Depletion of *npm1a* led to a decrease in both senescent and immune cell numbers, as well as decreased MG-derived proliferation in the retina. Together, the data support an unappreciated role for *npm1a* that promotes regeneration through both modulation of MG-derived proliferation and regulation of senescent cell function. Coupled with its known interaction with the MG-derived progenitor gene *chx10*/*vsx2*, defining the precise role of *npm1a* will be important to determine whether it might be a therapeutically relevant target in the retina after damage.

### SASP Heterogeneity and Cell-type Expression

After light damage, we detected early enrichment of SASP factors, led primarily by *mmp9*, *npm1a*, and *anxa2a* (Kramer, Gurdziel et al. 2021). Elevated expression of these factors persisted through 5 dpi, but after 10 dpi and out to 28 dpi, their levels returned to baseline, consistent with previous findings that the senescence response is transient and that senescent cells are cleared as regeneration is completed around 21-28 dpi (Konar, Flickinger et al. 2024). For NMDA damage, we observed upregulation of specific SASP factors at 3 dpi after damage, but expression of *txnb*, *jun*, and *anxa2a* did not return to baseline even out to 20 dpi. This could be attributed to the fact that these genes are expressed in resting MG or cell types that are not directly involved in regeneration, such as pericytes. Overall, upregulated SASP factors after NMDA damage remained elevated longer than observed following light damages, which may illustrate key differences in the timelines for repair between the two damage models.

Chronic light damage models showed less up-regulation of SASP factors as compared to acute light damage models. Chronic damage is an ongoing process, and cells are often in a heterogeneous state of degeneration and regeneration. This likely means that fewer cells are in a senescent state and thus have a lower SASP factor expression overall. While a few RASS factors were expressed in the chronic light damage dataset, there were only 22 SASP factors upregulated in chronic light damage compared to over 131 factors in acute light damage. Altogether, this lower SASP factor expression in chronic damage suggests that the role of senescence and SASP is either different between chronic vs. acute damage modalities or the expression levels across the retina are too heterogeneous to detect significant differences.

The use of chronic genetic models allowed us to identify conserved SASP factors compared to acute damage. The *bbs2^-/-^* and *cep290^-/-^* mutants show progressive loss of photoreceptors, either by 4 mpf (*bbs2*) or 6 mpf (*cep290*), highlighting a difference in the speed of degeneration in early adulthood (Grabinski, Parsana et al. 2023). The *bbs2^-/-^* mutants showed a much more pronounced SASP phenotype when compared to controls, than did the *cep290^-/-^* mutants. Despite differences in the speed and nature of photoreceptor degradation, we still observed many similar SASP factors and senescence markers enriched between the two mutants (*stat1b*, *cxcl8a*, *cd44a*, *txnb*, *jun*), including several SASP factors that were identified using acute damage models (*mmp9, anxa2a*). Since these zebrafish were collected at 6 mpf, the *bbs2^-/-^* mutant was past peak photoreceptor degeneration and the *cep290^-/-^* mutant was actively undergoing photoreceptor degeneration. This timing could explain why *bbs2^-/-^* mutants had a more pronounced SASP, as SASP factors associated with regeneration may have been more active than in the *cep290^-/-^* mutants.

The Regeneration-associated Senescence Signature (RASS) highlights a novel set of genes that are specific to both senescence and regeneration. This represents an important advancement in the understanding of how senescence modulates the retinal microenvironment and helps to facilitate proper regeneration. Of the 29 conserved SASP factors, only 3 (*mmp9, ahcy, hmga1a*) have been directly shown to influence retina regeneration, though many others have known roles in the regeneration of other tissues (Hoang, Wang et al. 2020, Silva, Nagashima et al. 2020, Campbell, El-Hodiri et al. 2023). The functions of the other 26 factors include chromatin modifiers, inflammation regulators, ECM remodelers, and factors that influence stemness, all of which are consistent with changing gene expression patterns during regeneration. By further studying other members of the RASS, this represents a promising avenue for the discovery of other pro-regenerative factors that originate from the SASP of DISI cells.

### SASP, Stem Cells, and Regeneration

The zebrafish retina responds to damage through MG, which produce progenitor cells that can differentiate and replace any lost cell type (Wan and Goldman 2016). For MG to facilitate this process, the local microenvironment must undergo dynamic changes to modulate signals needed for progenitor cell formation and the progression of regeneration (Lust and Wittbrodt 2018). This involves the activation of many signaling pathways including MAPK and PI3K, activation of transcription factors such as *ascl1a*, and the presence of growth factors such as *lin28* (Gallina, Todd et al. 2013, Gorsuch and Hyde 2013, Konar, Ferguson et al. 2020). The microenvironment is also regulated by the inflammatory state, including through NF-κB-mediated transcription of pro-inflammatory cytokines (Palazzo, Todd et al. 2022, Bludau, Weber et al. 2024, Perkins, Parsana et al. 2024). The SASP response to damage is a key facet in regulation of the microenvironment. Analysis of the RASS revealed an elevated expression of inflammatory factors and secreted factors that activate genes that promote wound healing, pluripotency, and growth (Coppé, Desprez et al. 2010, Ritschka, Storer et al. 2017, Saul, Kosinsky et al. 2022). In the zebrafish retina, premature loss of senescent cells truncates retina regeneration, indicating the influence these factors exert on the activation state of the MG after damage (Konar, Flickinger et al. 2024). Several enriched SASP factors showed a common function of regulating stemness, including in MG. While it is thought that an appreciable number of senescent cells are immune-derived after damage, both the bulk RNAseq and scRNAseq datasets indicate that SASP factors are also expressed and presumably released by glial cells, including MG (Acosta, Banito et al. 2013, Ring, Valdivieso et al. 2022). This highlights the heterogeneity of SASP expression in the retina, as across the full list of over 700 factors in the SASP atlas, only 29 showed common upregulation, and they were split between MG and immune-derived cells. Overall, SASP expression during retina regeneration is a dynamic process that likely plays a key role in both modulating inflammation and generating a highly favorable pro-regenerative microenvironment for MG to initiate regeneration, produce progenitor cells, and for those cells to differentiate into lost cell types as regeneration proceeds.

### Conclusion

The detection of senescent cells supports a model in which the secretion of SASP factors during retina regeneration provides crucial microenvironmental cues that can induce MG to initiate regeneration, produce progenitor cells, and promote regeneration. Across multiple acute and chronic retina damage models, we discovered a novel Regeneration-associated Senescence Signature (RASS) that includes 29 SASP factors and 2 senescence markers commonly upregulated after damage. These factors modulate inflammation, chromatin dynamics, and stemness, which together combine to initiate and allow progression of regeneration. Identification of SASP factors and understanding the dynamic SASP response in an innately regenerative model organism will help to identify candidate factors that promote regeneration in organisms incapable of spontaneous retina regeneration.

## Acknowledgements

The authors would like to thank Jack Hollander for his assistance with maintaining the zebrafish aquatic facility throughout the experimentation process. This work was supported by Vanderbilt University and the Stevenson family endowment.

