## Supplemental Figures 1-6 for "Analysis of The Senescence Secretome During Zebrafish Retina Regeneration"

**Supplemental Figures for Konar et al, 2025.**

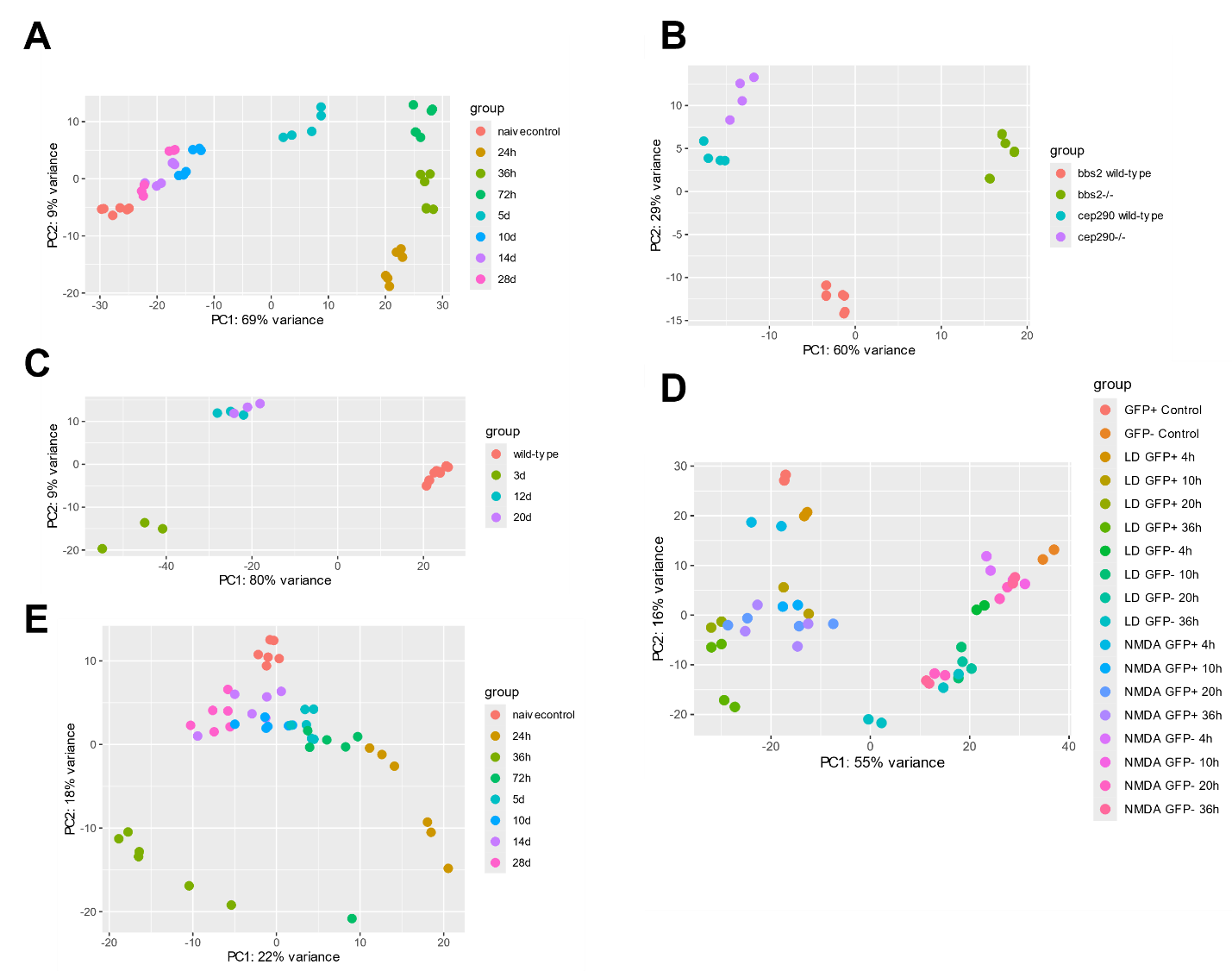

**Supplemental Figure 1. Principal component analysis of bulk RNAseq datasets.** A) Kramer et al. 2021, B) Grabinski et al. 2023, C) Konar et al. 2025 (this publication), D) Mitchell et al. 2019, E) Hoang et al. 2020, F) Kramer et al. 2023.

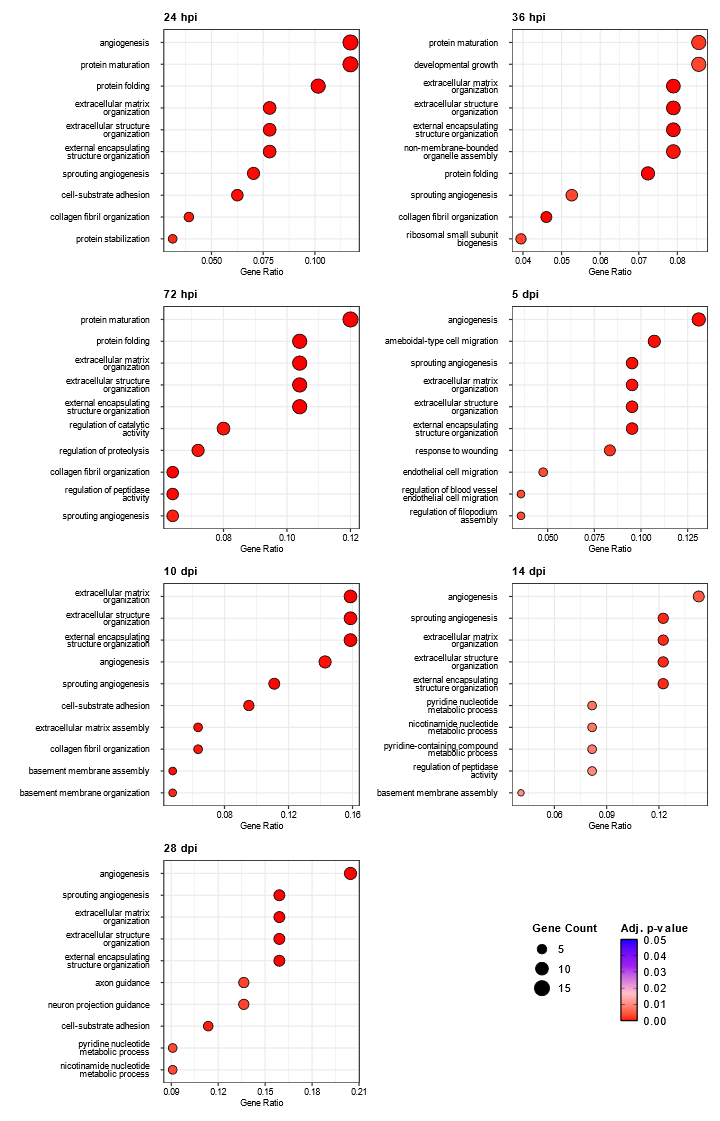

**Supplemental Figure 2. Biological process Gene Ontology for enriched SASP factors after acute light damage.** Bulk RNAseq was performed on whole zebrafish retinas after acute light damage and compared to undamaged controls. For each time point, Gene Ontology enrichment for biological process was conducted for all SASP factors and markers enriched beyond a log2FC of 1 with adjusted p-values < 0.05 using clusterProfiler. The top 10 enriched terms for the respective time points are displayed as dot plots.

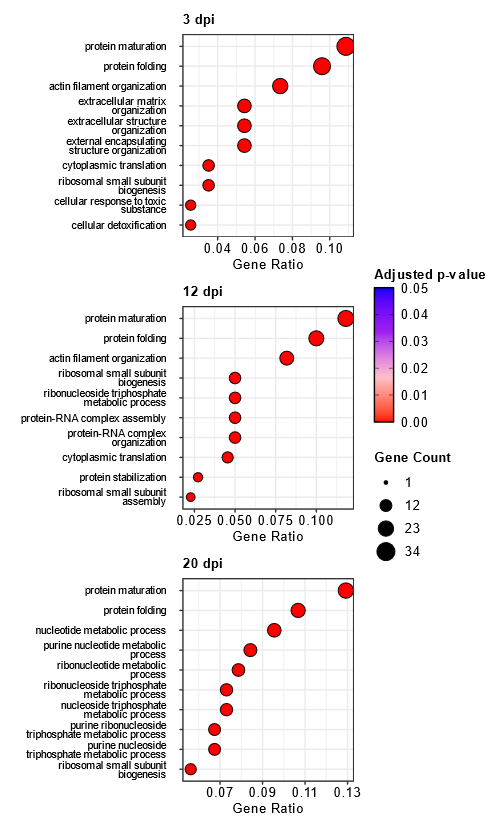

**Supplemental Figure 3. Biological process Gene Ontology for enriched SASP factors after NMDA damage.** Bulk RNAseq was performed on whole zebrafish retinas after intravitreal injections of NMDA and compared undamaged controls (Grabinski, Parsana et al. 2023). For each time point, Gene Ontology enrichment for biological process was conducted for all SASP factors and markers enriched beyond a log2FC of 1 with adjusted p-values < 0.05 using clusterProfiler. The top 10 enriched terms for the respective time points are displayed as dot plots.

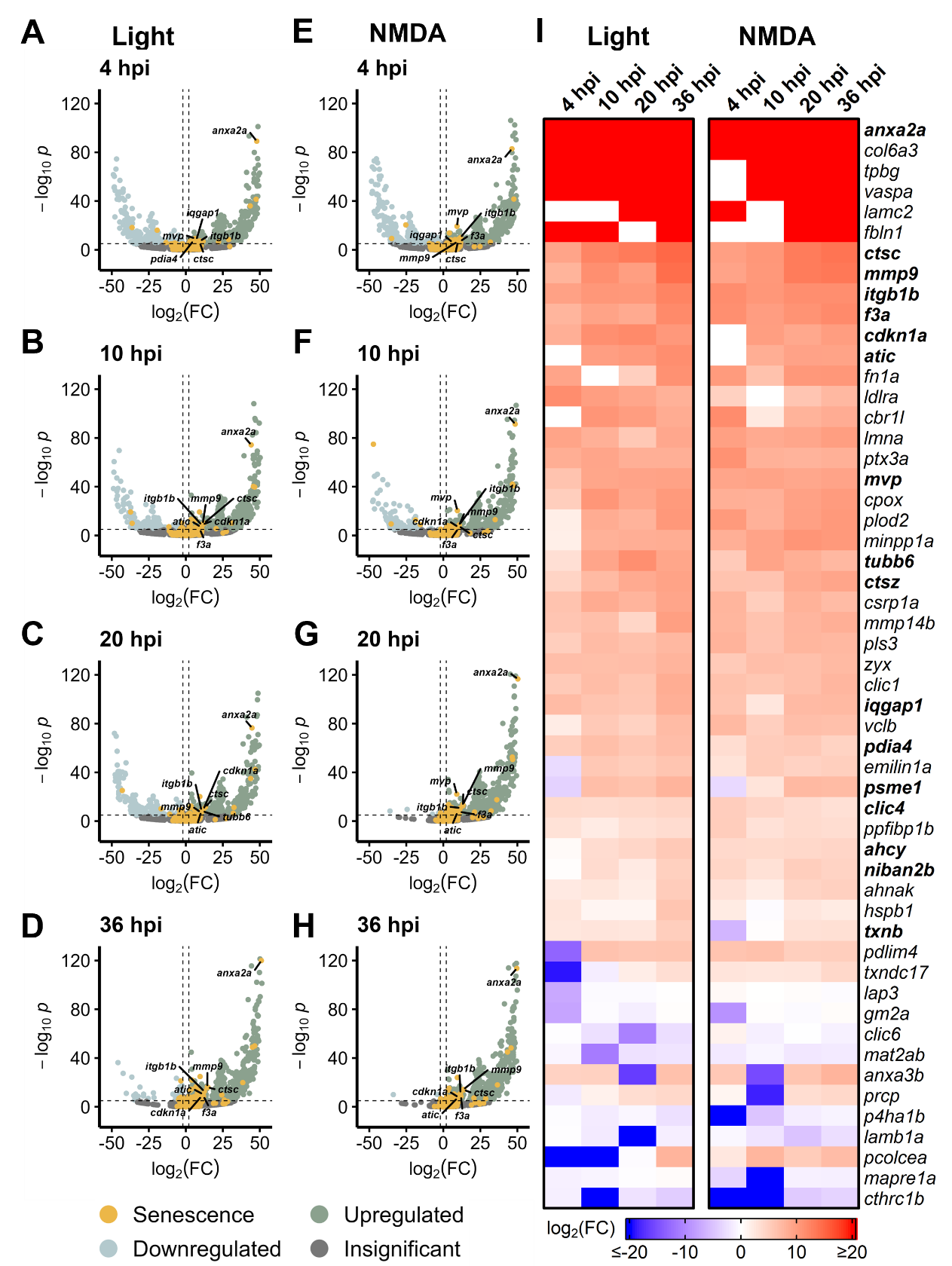

**Supplemental Figure 4. SASP factor conservation across acute damage models in non-Müller glia.** Bulk RNAseq was performed on non-Müller glia after either NMDA or light damage compared to undamaged controls. (A-D) Volcano plots showing differentially expressed SASP factors at the indicated times post light damage. (E-H) Volcano plots showing differentially expressed SASP factors at the indicated times post NMDA damage. Dashed lines represent values with a |log2FC| > 2 and p-values < 10-6. Select up-regulated factors are labeled for each time point. (I) Heatmap showing enrichment of SASP factors at the indicated times post light or NMDA damage. SASP factors were included in the heatmap if they had a log2FC > 5 with p-values < 10-6 for at least one time point; bolded genes are part of the RASS.

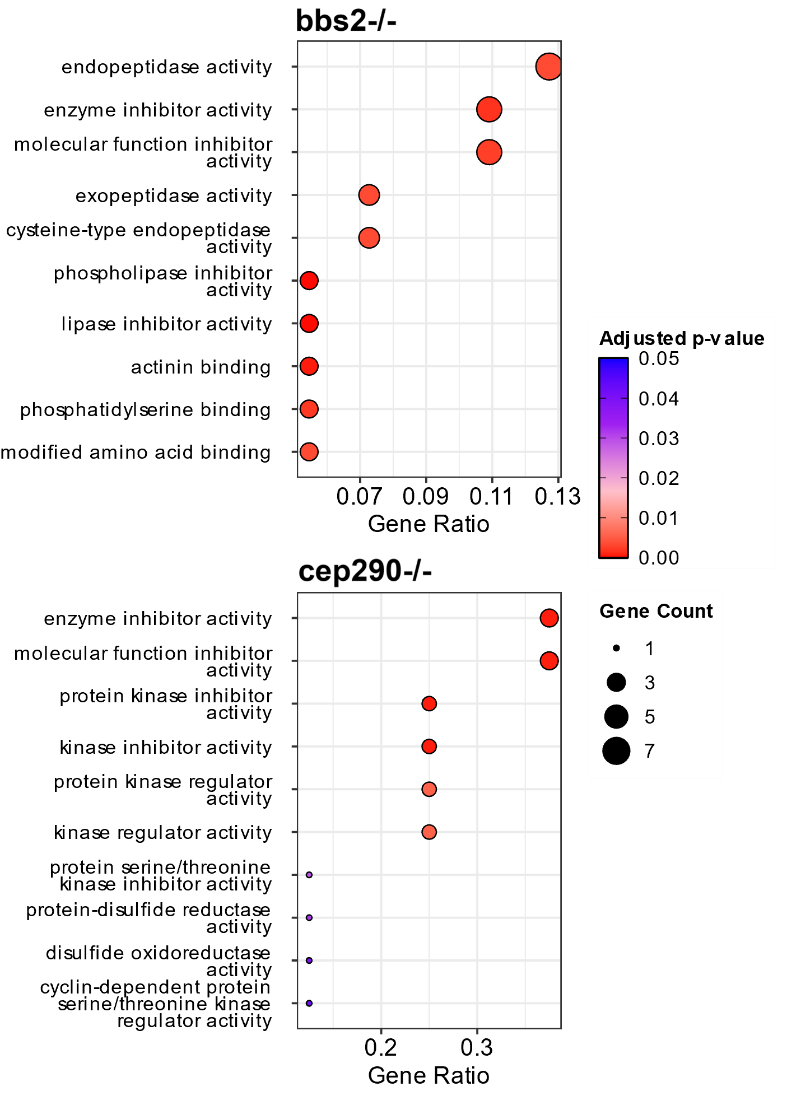

**Supplemental Figure 5. Molecular Function Gene Ontology for enriched SASP factors in genetic mutants with chronic photoreceptor damage.** Bulk RNAseq was performed on whole zebrafish retinas from bbs2-/- and cep290-/- ciliopathy mutants. For each genetic mutant, Gene Ontology enrichment for molecular function was conducted for all SASP factors and markers enriched beyond a log2FC of 1 with adjusted p-values < 0.05 using clusterProfiler. The top 10 enriched terms for the respective time points are displayed as dot plots.

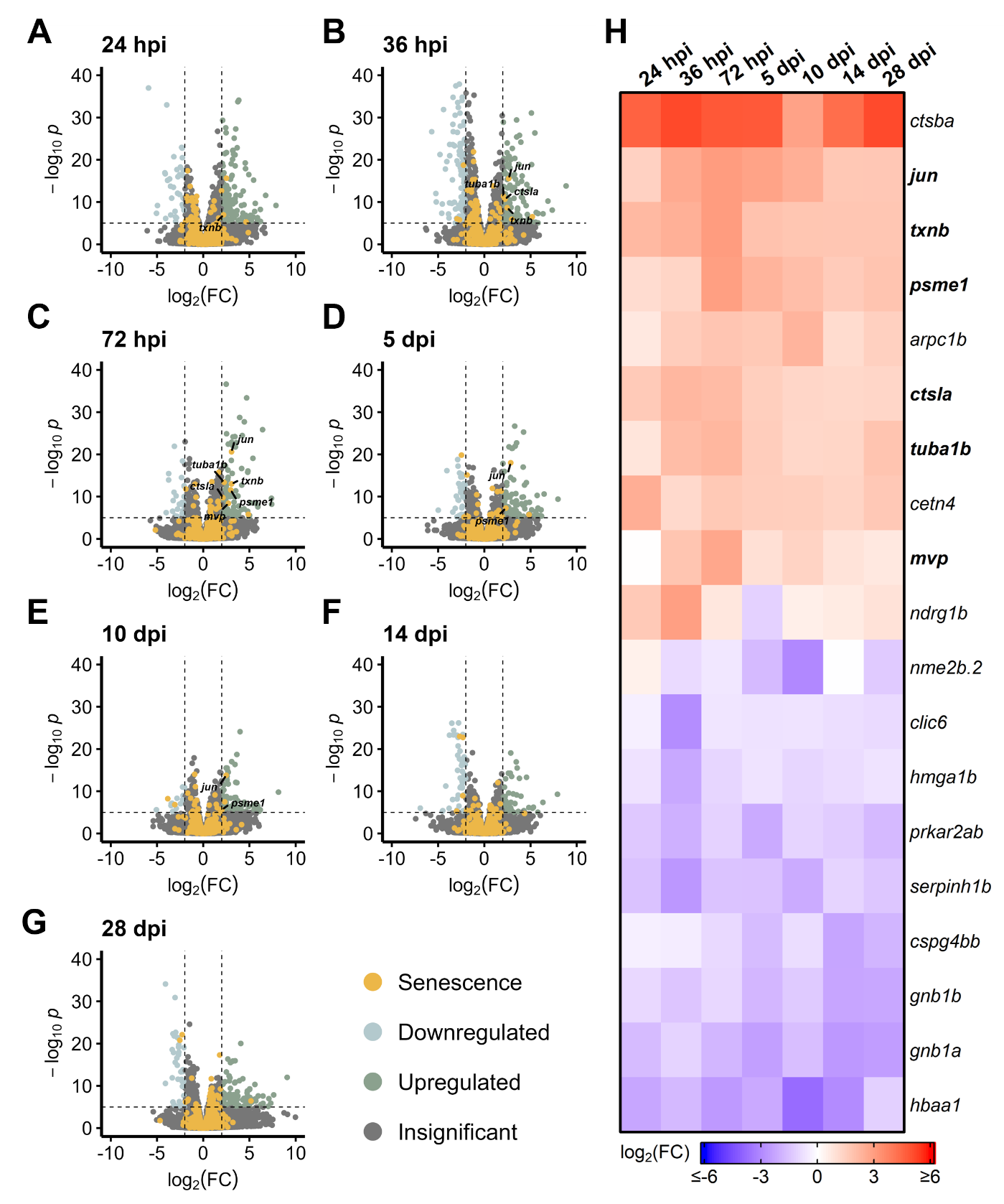

**Supplemental Figure 6. Differentially expressed SASP factors after chronic light damage.** Bulk RNAseq was performed on whole zebrafish retinas after chronic light damage. (A-G) Volcano plots depicting up- and down-regulated SASP factors and senescence markers at the indicated times post damage. Dashed lines represent values with a |log2FC| > 2 and p-values < 10-6. Select factors are labeled for each time point. (H) Heatmap showing enrichment of SASP factors at the indicated times post chronic light damage. SASP factors and senescence markers were included in the heatmap if they had a log2FC > 2 with p-values < 10-6 for at least one time point; bolded genes are part of the RASS.
